# Neurophysiological Substrates of Configural Face Perception in Schizotypy

**DOI:** 10.1101/668400

**Authors:** Sangtae Ahn, Caroline Lustenberger, L. Fredrik Jarskog, Flavio Fröhlich

**Affiliations:** Department of Psychiatry, University of North Carolina at Chapel Hill, Chapel Hill NC 27599; Carolina Center for Neurostimulation, University of North Carolina at Chapel Hill, Chapel Hill NC 27599; Mobile Health Systems Lab, Institute of Robotics and Intelligent Systems, ETH Zurich, 8092 Zurich, Switzerland; North Carolina Psychiatric Research Center, Raleigh, NC, 27610; Department of Neurology, University of North Carolina at Chapel Hill, Chapel Hill NC 27599; Department of Biomedical Engineering, University of North Carolina at Chapel Hill, Chapel Hill NC 27599; Department of Cell Biology and Physiology, University of North Carolina at Chapel Hill, Chapel Hill NC 27599; Neuroscience Center, University of North Carolina at Chapel Hill, Chapel Hill NC 27599

**Keywords:** face perception, schizotypy, electroencephalography, event-related potential

## Abstract

Face perception is a highly developed function of the human visual system. Previous studies of event-related potentials (ERPs) have identified a face-selective ERP component (negative peak at about 170 milliseconds after stimulation onset, N170) in healthy participants. In contrast, patients with schizophrenia exhibit reduced amplitude of the N170, which may represent a pathological deficit in the neurophysiology of face perception. Interestingly, healthy humans with schizophrenia-like experiences (schizotypy) also exhibit abnormal processing of face perception. Yet, it has remained unknown how schizotypy in healthy humans is associated with the neurophysiological substrate of face perception. Here, we recruited 35 participants and assessed their schizotypy by the magical ideation rating scale. We used high-density electroencephalography to obtain ERPs elicited by a set of Mooney faces (face and non-face conditions). We divided the participants into two groups (high and low schizotypy) by a median split of schizotypy scores. We investigated mean reaction times and the N170 component in response to the stimuli. We found significant slowed reaction times and reduced amplitude of the N170 component in response to the face stimuli in the high-schizotypy group. In addition, across the full data set, we found that the schizotypy scores were significantly correlated with both the reaction times and the N170 amplitude. Our results thus support the model of schizotypy as a manifestation of a continuum between healthy individuals and patients with schizophrenia, where the N170 impairment serves as a biomarker for the degree of pathology along this continuum.

## Introduction

Face perception is a highly developed function of the human visual system (Haxby et al., 2000). Face perception develops at a very early age; faces are the first distinguishable object for infants (Morton and Johnson, 1991). Previous functional brain imaging studies have observed that several brain regions are involved in face perception including the fusiform face area (Kanwisher et al., 1997), right lateral occipital area (Gauthier et al., 2000), and superior temporal sulcus (Hoffman and Haxby, 2000). Complementing the high spatial resolution of imaging, electrophysiological recordings with high temporal resolution provide insights into the neuronal network dynamics of face perception (Rossion, 2014). In particular, event-related potentials (ERPs), which represent electrical potentials elicited by time-locked external stimuli, can be measured with electroencephalography (EEG) and have been widely investigated in the study of the neural basis of face perception. The N170 component, which is a negative peak potential at approximately 170ms from the stimulus onset, is considered a face-selective ERP component (Bentin et al., 1996; Jeffreys, 1996; Rossion and Jacques, 2011). A greater increase in amplitude of the N170 components was observed for face stimuli when compared to other objects in healthy participants (Rossion and Caharel, 2011). In contrast, patients with schizophrenia exhibit impaired processing of face perception (Kohler et al., 2010; McCleery et al., 2015; Whittaker et al., 2001). In particular, they showed reduced amplitude (Herrmann et al., 2004; Ibáñez et al., 2012; Lynn and Salisbury, 2008; Turetsky et al., 2007) and delayed latency (Caharel et al., 2007) of the N170 component in response to face stimuli along with structural deficits in bilateral anterior and posterior fusiform gyrus gray matter volumes (Onitsuka et al., 2006). Importantly, these deficits may represent symptoms severity of schizophrenia (Akbarfahimi et al., 2013; Campanella et al., 2006; Kim et al., 2013; Maher et al., 2016; Zheng et al., 2016).

Schizotypy refers to a latent personality trait, which is prone to schizophrenia (Meehl, 1962). Previous studies have shown that individuals with schizotypy appear to share schizophrenia risk genes (Baron and Risch, 1987; Cadenhead and Braff, 2002; Vollema et al., 2002). Several neuroimaging studies found structural (Moorhead et al., 2009; Peters et al., 2010) and functional (Lagioia et al., 2010; Volpe et al., 2008) impairment in participants with schizotypy, which are also found in patients with schizophrenia. These phenomena can be explained by a theory of a continuum between patients with schizophrenia and healthy participants (Cochrane et al., 2012; van Os et al., 2009). Interestingly, participants with schizotypy exhibit impaired processing of face perception in behavioral tasks (Dickey et al., 2011; Mikhailova et al., 1996; Platek and Gallup, 2002). Yet, the neurophysiological substrate of configural face perception, which refers to the ability in recognizing whether a stimulus is a face or not, associated with schizotypy has remained unknown. Here, we recruited healthy participants and assessed their magical ideation. Magical ideation was chosen as an indicator of schizotypy (Eckblad and Chapman, 1983) since it has been considered as a prominent symptom of schizotypy (Karcher and Shean, 2012; Kwapil et al., 1997; Thalbourne, 1994) We hypothesized that participants with higher schizotypy exhibit more impaired processing of faces both in terms of behavioral performance and neurophysiological correlates. We investigated reaction times and the face-selective N170 components in response to face stimuli and its relationship with the schizotypy scores. Our findings suggest that deficits of configural face perception in healthy humans with high schizotypy represent a continuum with schizophrenia and the neurophysiological substrate may represent the degree of this continuum.

## Methods

### Study design

This study was performed at the University of North Carolina at Chapel Hill (UNC-CH) and approved by the Biomedical Institutional Review Board of UNC-CH. Participants were recruited from UNC-CH community. All participants were males, right-handed, and free of any neurologic or sleep disorders (age: 21.8±3.5 years, mean±std). A urine drug screen was performed to exclude participants who tested positive for drugs of abuse. Eligibility of participants was determined by a telephone screening. All participants provided written informed consent form before participation. Thirty-six participants were enrolled in the study. Participants completed questionnaires for inclusion/exclusion criteria, handedness, and schizotypal traits (schizotypy). The Edinburgh handedness inventory was used and a laterality index was obtained from each participant. Data from 35 participants were analyzed since one participant did not complete the schizotypy questionnaire.

### Schizotypy

We used an adapted version of the magical ideation scale (Eckblad and Chapman, 1983) as an indicator of schizotypal traits (schizotypy). This assessment originally consisted of 30 true/false questions and we had previously adjusted the questions with a six-point rating scale (0: strongly disagree, 5: strongly agree) for a fine-tuned assessment of schizotypal traits (Lustenberger et al., 2015). Summation of the scores for the 30 questions were used as a schizotypy score (0: no schizotypy, 150: high schizotypy). Participants were divided into two groups – low and high schizotypy – using a median split. Schizotypy scores and other questionnaire scores, including demographics for these groups, are reported in Table 1.

**Table 1.**
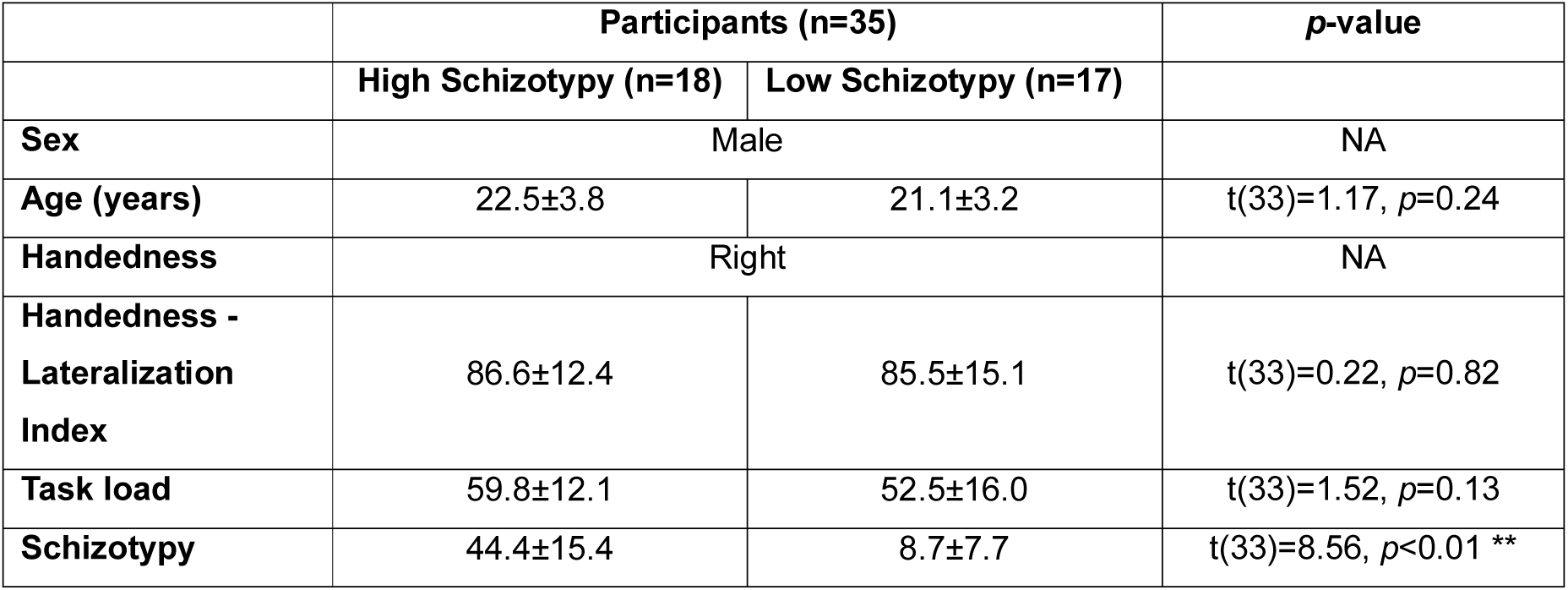
Demographics and schizotypy scores. Mean and standard deviation are presented. Two sample t-test was used for statistical tests. \*\**p*<0.01

|  | Participants (n=35) |  | p-value |
| --- | --- | --- | --- |
|  | High Schizotypy (n=18) | Low Schizotypy (n=17) |  |
| Sex | Male |  | NA |
| Age (years) | 22.5±3.8 | 21.1±3.2 | t(33)=1.17, $p=0.24$ |
| Handedness | Right |  | NA |
| Handedness - Lateralization Index | 86.6±12.4 | 85.5±15.1 | t(33)=0.22, $p=0.82$ |
| Task load | 59.8±12.1 | 52.5±16.0 | t(33)=1.52, $p=0.13$ |
| Schizotypy | 44.4±15.4 | 8.7±7.7 | t(33)=8.56, $p < 0.01$ ** |

### Stimuli and task

We used 80 Mooney faces, which refer to two-tone pictures of faces, that have the highest face-like ratings from a validated set consisting of 144 Mooney faces (Verhallen and Mollon, 2016). We inverted and scrambled the 80 Mooney faces to make another set of 80 non-face stimuli (Uhlhaas et al., 2006). All non-face stimuli had same contrast (white/black) with the face stimuli. The size of all stimuli was 6.8 x 10 cm (width x height). Participants performed a face perception task consisting of the 80 face and 80 non-face stimuli. First, written task instructions were displayed for 60 seconds and randomized face stimuli were presented for 200 milliseconds each. A total of 160 trials were presented (80 for face and 80 for non-face stimuli) with an inter-trial interval of 3.5 to 4.5 seconds. Participants were randomized to one of two task forms (task A and task B) in an equal distribution. In task A, participants were asked to press the left-arrow key for face stimuli and right-arrow key for non-face stimuli as precisely and quickly as possible. In task B, the assignment of the two response keys was reversed (Figure 1). All instructions and experimental tasks were implemented in Presentation (Neurobehavioral Systems Inc., Berkeley, CA). After the task, participants completed a task load questionnaire, which assess how much the task is difficult to participants. EEG data were recorded by a high-density EEG system (128 channels, EGI Inc., Eugene, OR) at a sampling rate of 1kHz. Channel Cz and one channel between Cz and Pz were used as the reference and ground, respectively.

**Figure 1.**
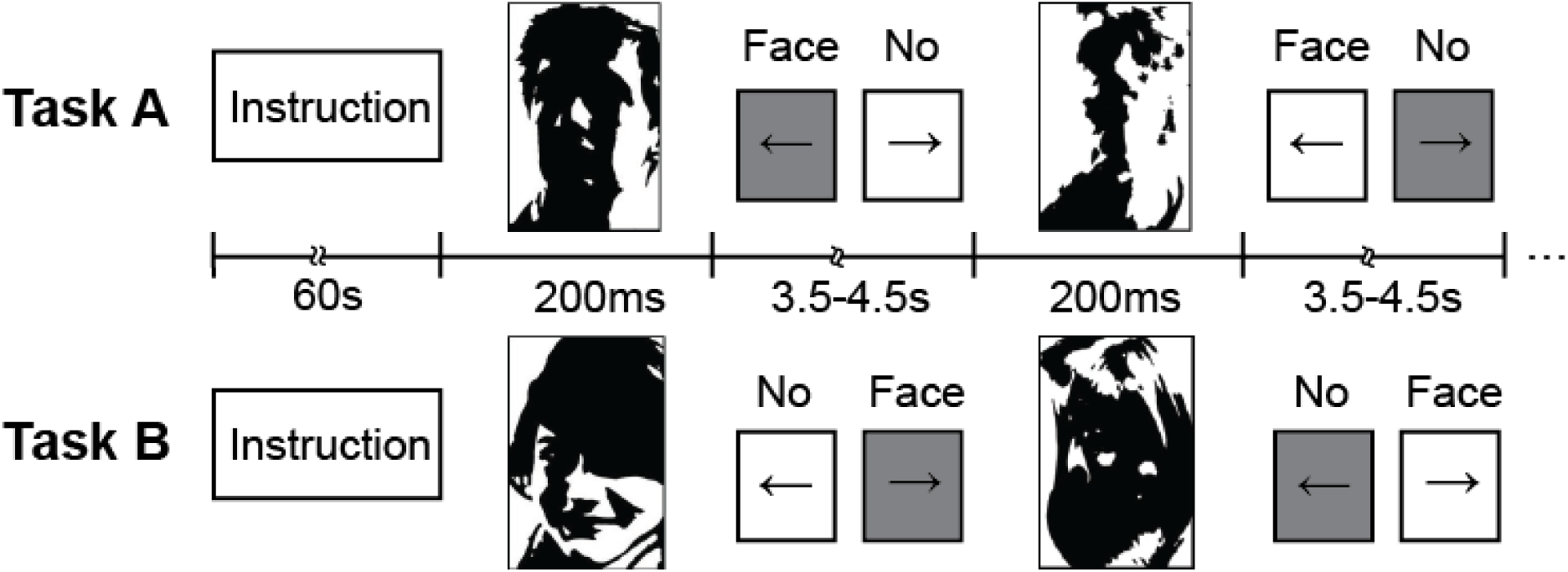
Experimental paradigm for configural face perception using a set of Mooney faces. Instruction was displayed for 60 seconds and then face or non-face stimuli were presented for 200 milliseconds. Participants were asked to press the left or right arrow key according to the type of task as quickly and precisely as possible. Inter-trial interval was 3.5-4.5 seconds. The type of key press was randomized and counter-balanced across participants (task A and task B). A total of 160 trials (80 for each stimulus condition) were obtained.

### Analysis and statistical testing

Reaction times were calculated from the key press in response to the face and non-face stimuli. Outliers (>2 seconds) and missed trials were removed and mean reaction times were obtained for each participant. For EEG data, offline processing was performed by EEGLAB (Delorme and Makeig, 2004), FieldTrip (Oostenveld et al., 2010), and custom-built scripts in MATLAB R2015b (Mathworks, Natick, MA). First, all data were band-pass filtered from 1 to 50Hz and then downsampled to 500Hz. Second, the data were preprocessed by an artifact subspace reconstruction algorithm (Mullen et al., 2013) to remove high-variance signals at each channel and identify noisy channels. Third, noisy channels that were found in the previous step were interpolated and common average referencing was performed. Lastly, infomax independent component analysis (ICA) (Jung et al., 2000) was performed to remove eye blinking, eye movement, muscle activity, and heartbeats artifact. All ICA components were visually inspected and components were manually selected for rejection. After the preprocessing steps, data were epoched from −100ms to 600ms according to the stimuli onset. A total of 160 trials (80 for face and 80 for non-face stimuli) were obtained for each participant. Each trial was visually inspected in the time domain and noisy and missed trials were removed. Then, the remaining trials were averaged at each channel for each participant to obtain ERPs.

For statistical testing, we used the two sample (un-paired) t-test for demographics including age, handedness lateralization index, task load, and schizotypy scores (Table 1) between the high and low schizotypy groups. In behavioral and neurophysiological data, we used a linear mixed-effects model analysis with fixed factors of “group” (high and low schizotypy), “task type” (task A and task B), and “condition” (face and non-face), with a random factor of “participant” written in R (R foundation for Statistical Computing, Vienna, Austria). The dependent variables were mean reaction times and amplitude of the N170 components. Post-hoc statistical tests were performed using two sample t-tests between the two groups. We used the Spearman’s rho for calculating correlations since the schizotypy scores are ordinal.

To calculate statistical significance for ERPs, we adopted a non-parametric cluster-based permutation test (Maris and Oostenveld, 2007) to deal with the multiple comparison problem of high-density EEG. First, t-tests were conducted for each channel and time point across participants between conditions (e.g. face vs. non-face). We then constructed clusters from obtained spatio-temporal significant t-value map (*p*<0.05) and summed all the positive or negative t-values within the clusters separately. The significant t-values were clustered based on spatio-temporal adjacency. The minimum size of a cluster was set to two points. A neighboring channel was defined as spatial adjacency within 4 cm (Maris and Oostenveld, 2007). For permutation test, we shuffled all trials and divided it into two datasets. We then conducted t-tests for the two datasets to obtain a t-value map. We repeated this procedure by the Monte Carlo simulation with 1000 iterations. To compare with the original dataset, we extracted the largest cluster from each permutation test. Lastly, we constructed a histogram of the 1000 values of the cluster-level statistics and calculated a probability density function (PDF) to estimate cluster-level *p*-values. The input for the PDF was the cluster-level statistics from the original dataset, while the output was a *p*-value for each cluster-level statistic. The cluster-level *p*-values were corrected and approximated by this cluster-based permutation test.

### Inter-stimulus perceptual variance

One study (Thierry et al., 2007) claimed that inter-stimulus perceptual variance (ISPV) may affect amplitude in the N170 component. To investigate ISPV in stimulus images used in our study, we first chose images (N=1, 10, 20, 40, and 80) randomly from the image set for each stimulus condition (face and non-face) and computed pixel-by-pixel averaging across the stimulus images. We then computed histogram of pixel-by-pixel correlations between the stimulus images for each condition to quantify ISPV. Non-parametric kernel-smoothing.was used to estimate the probability density function of the histogram.

## Results

### Behavioral data

The schizotypy scores between the two groups, which were determined using a median split, were significantly different (two sample t-test, high schizotypy: n = 18, low schizotypy: n=17; t(33)=8.56, *p*<0.01, Table 1). We then investigated mean reaction times for the task and found a significant interaction between “condition” and “group” (F_1,31_=5.7, *p*=0.02) using a linear mixed-effects model. A post-hoc two sample t-test (high vs. low schizotypy) yielded a significant difference only in the face condition (t(33)=2.35, *p*=0.02, Table 2, Figure 2A). Hit rate in both face and non-face conditions was not significantly different between the two groups (F_1,31_=0.22, *p*=0.64, Table. 2). We next calculated the Spearman’s rho between the mean reaction times and schizotypy scores across all participants. We found a significant positive correlation (rho=0.47, *p*=0.004), which indicates that button responses were slower as participants had higher schizotypy scores (Figure 2B). These findings show that the high schizotypy group exhibited delayed responses in a face perception task and that the delay was significantly correlated to the degree of schizotypy scores.

**Table 2.** Behavioral and neurophysiological data. Mean and standard deviation are presented. A mixed-effects model analysis and two sample t-test were used for statistical tests. \**p*<0.05, \*\**p*<0.01

|  | Participants (n=35) |  | p-value |
| --- | --- | --- | --- |
|  | High Schizotypy (n=18) | Low Schizotypy (n=17) |  |
| **Reaction time (ms)** |  |  |  |
| **Face** | 390.8±33.8 | 343.3±22.4 | $t(33)=2.35$, $p=0.02$ * |
| **Non-Face** | 357.1±44.1 | 351.8±33.7 | $t(33)=0.57$, $p=0.57$ |
| **Hit rate (%)** |  |  |  |
| **Face** | 74.4±4.4 | 75.2±5.2 | $F_{1,31}=0.22$, $p=0.64$ |
| **Non-Face** | 76.8±5.8 | 78.1±7.1 |  |
| **Absolute N170 (uV)** |  |  |  |
| **Face** | 1.9±0.5 | 3.3±0.8 | $t(33)=-5.69$, $p<0.01$** |
| **Non-Face** | 2.4±0.3 | 2.3±0.4 | $t(33)=0.12$, $p=0.45$ |

**Figure 2.**
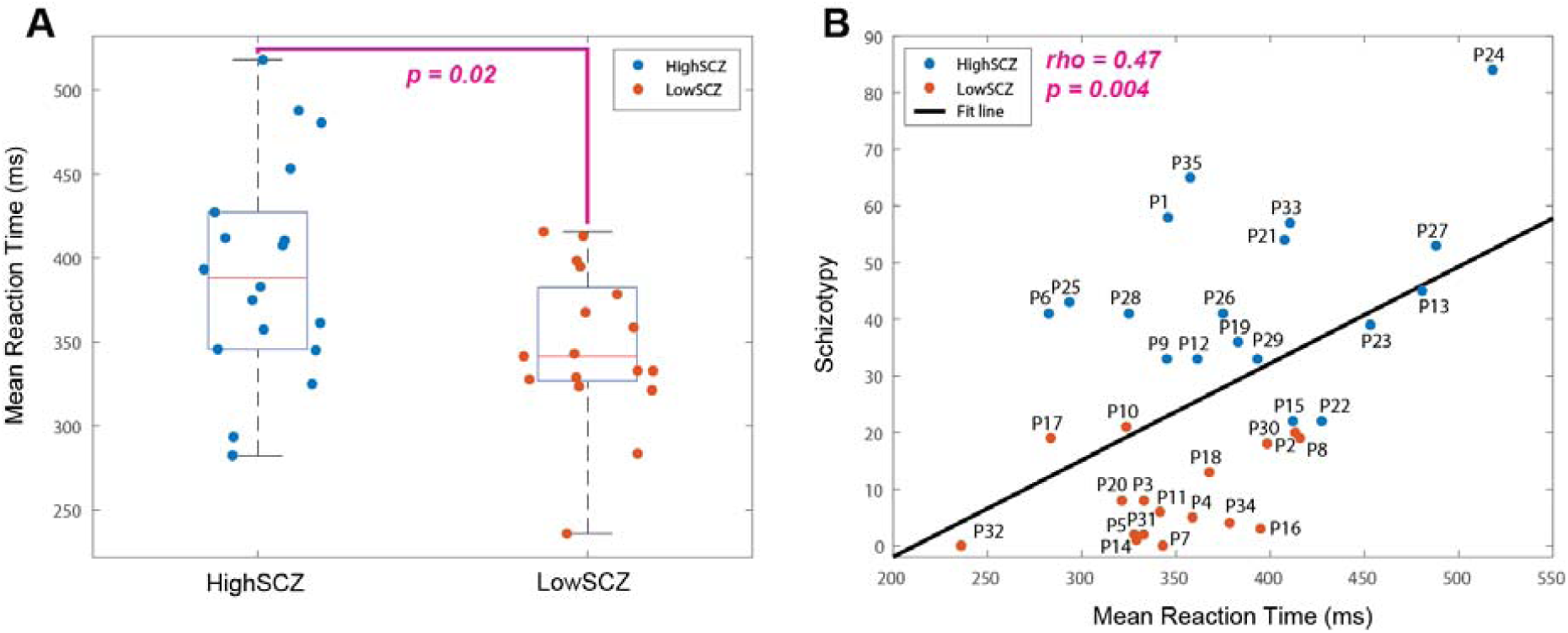
Mean reaction time to the face stimuli. (A) Boxplot on scatter plot of all participants (*p*=0.02). Blue and orange dots indicate participants from high-schizotypy (HighSCZ) and low-schizotypy (LowSCZ) groups, respectively. (B) Scatter plot between mean reaction time and schizotypy scores (rho=0.47, *p*=0.004). Black line indicates the least-squares fit line to the scatter plot.

### ERPs for configural face perception

We investigated the ERPs elicited by face and non-face stimuli for all participants. We averaged ERPs across correct trials at each channel for each participant and then averaged these across participants to obtain grand-averaged ERPs. Grand-averaged ERPs were obtained for both face and non-face stimulus conditions. This approach resulted in a total of 128 averaged ERPs matching the number of EEG channels (upper panels in Figures 3A and 3B). Topographical distributions for two ERP components (P100 and N170) were calculated (lower parts in Figures 3A and 3B). We found positive potentials at 100ms over the occipital region (P100) and negative potentials at 170ms over the bi-occipito-temporal region (N170). Differences in ERPs averaged over the occipito-temporal region indicate that the N170 component was strongly presented for the face stimulus condition (Figure 3C, upper panel, shaded gray bar indicates statistically significant time period determined by the cluster-based non-parametric test). We obtained topographical distributions for averaged differences (Face – Non-Face) in P100 and N170 components and found statistically significant channels over the occipito-temporal region in the N170 component (*p*<0.05, corrected for multiple comparisons, significant channels marked by asterisks, 11 channels). These findings demonstrate that the face-selective N170 components over the occipito-temporal region were elicited by the configural face perception task using a set of Mooney faces.

**Figure 3.**
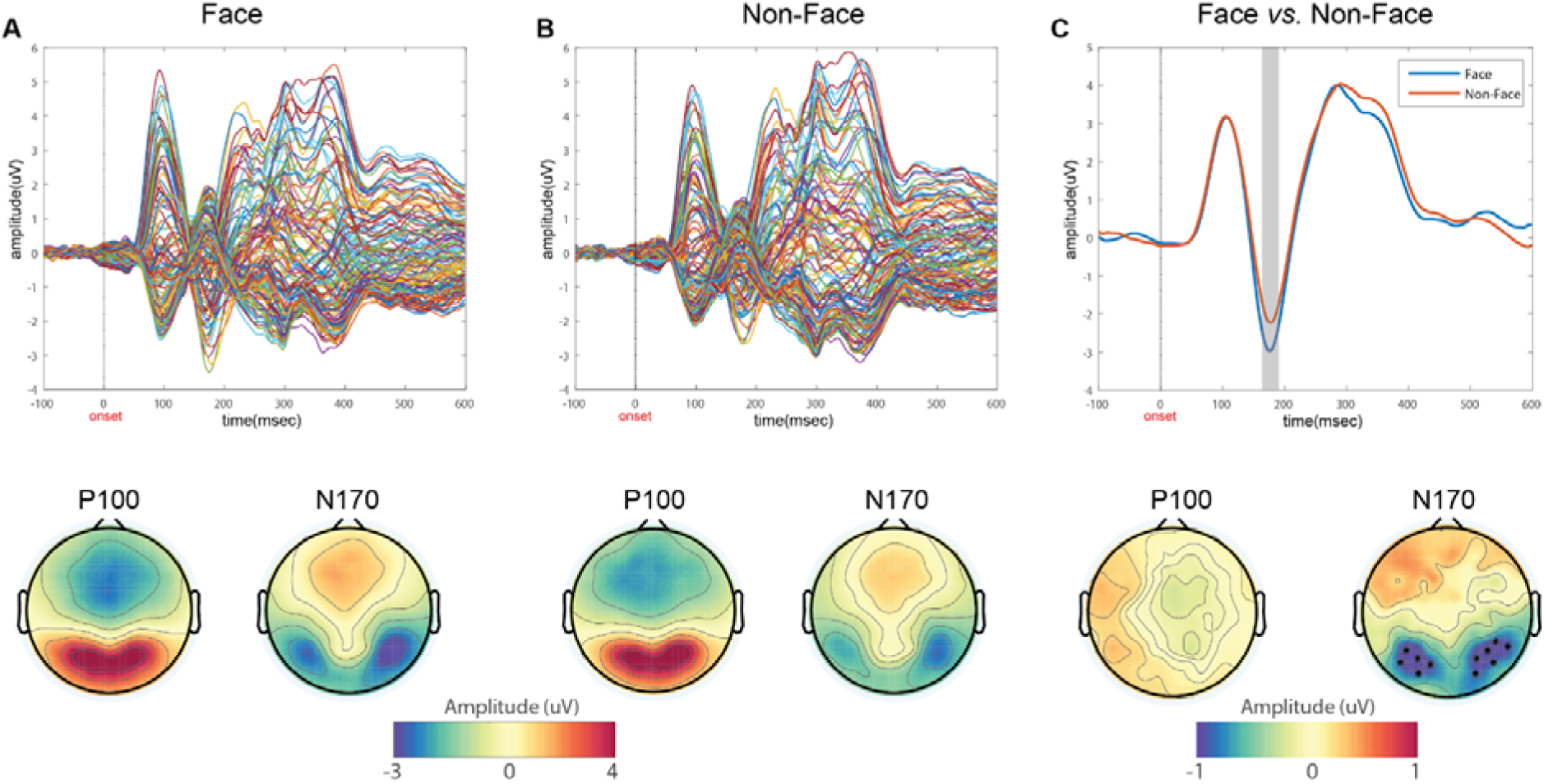
Grand-averaged ERPs for all channels and topographical distributions of ERP components (P100 and N170) for face and non-face conditions. ERPs and topographies for (A) face stimuli and (B) non-face stimuli. (C) Differences in ERPs averaged over the occipito-temporal region and topographies for P100 and N170. Asterisks in the topography represent statistically significant channels (*p*<0.05, corrected for multiple comparisons).

### ERPs between high- and low-schizotypy groups

To investigate ERP differences between the two groups, we calculated grand-averaged ERPs in both face and non-face conditions. ERPs over the occipito-temporal region were obtained for each group in the face and non-face condition. We found that the high-schizotypy group showed reduced amplitude of the N170 component in the face condition and differences between the two groups were statistically significant (Figure 4A, shaded gray bar, corrected). Topographical distributions indicate that the amplitude of the N170 component was reduced (absolute decrease) in the high-schizotypy group over the occipito-temporal region (Figure 4A, topographic map of N170, High – Low schizotypy). Marked asterisks represent statistically significant channels (*p*<0.05, corrected, 20 channels). No such difference was found for the P100 component. In contrast, we found no statistical significance for both P100 and N170 components in the non-face condition (Figure 4B). No channels were significantly different between the groups (Figure 4B, topographic maps). Further, we investigated the peak amplitude of the averaged N170 components across trials from all participants and found a significant interaction between “condition” and “group” (F_1,31_=31, *p*<0.01) using a linear mixed-effects model. A post-hoc t-test (high vs. low schizotypy) revealed a significant difference between two groups in the face condition (t(33)=-5.69, absolute amplitude, *p*<0.01, Table 2, Figure 4C). Strikingly, absolute peak amplitudes of the N170 components were significantly correlated with schizotypy scores (Figure 4D, rho=-0.56, *p*=0.0003), which indicates that great reduction in amplitude was associated with higher schizotypy scores. These findings show that the neurophysiological substrates of configural face perception is significantly reduced over the occipito-temporal region in the high schizotypy group in response to the face stimuli and the degree of reduction is significantly correlated with the schizotypy scores.

**Figure 4.**
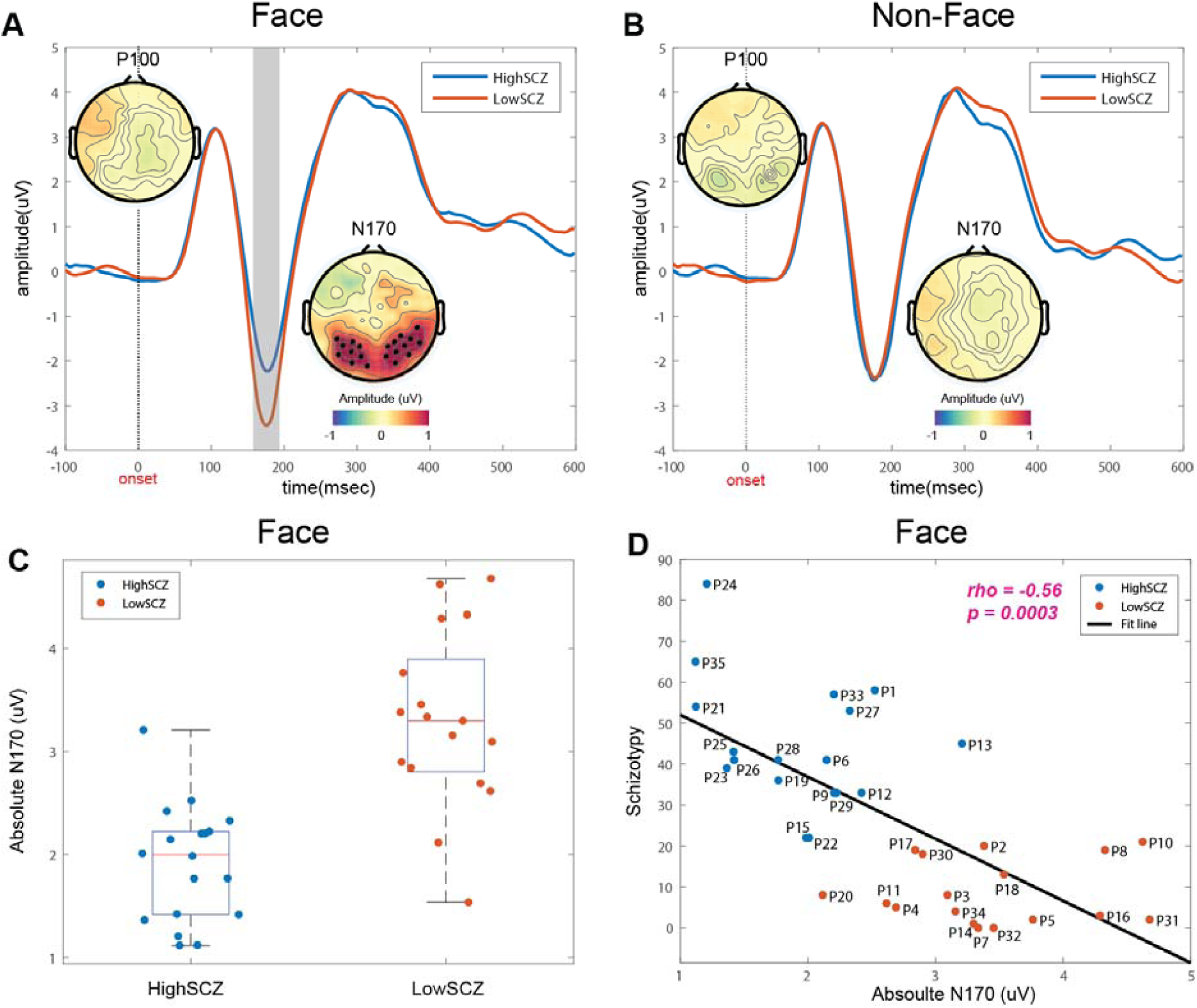
Grand-averaged ERPs for occpito-temporal channels and topographical distributions of ERP components (P100 and N170) in the two groups (high and low schizotypy, LowSCZ and HighSCZ) for the (A) face and (B) non-face conditions. Shaded gray bar in the ERP plot indicates significant time period and marked asterisks in the topography indicate significant channels (*p*<0.05, corrected). (C) Absolute N170 components between the high and low schizotypy groups in the face condition. (*p*<0.1) (D) Scatter plot of absolute N170 components and schizotypy scores (rho=-0.56, *p*=0.0003). Black line indicates the least-squares fit line to the scatter plot.

## Discussion

In this study, we asked how the neurophysiological substrate of configural face perception is associated with schizotypy in healthy participants. To answer this question, we adopted a configural face perception task consisting of a set of Mooney faces. We recorded high-density EEG data to obtain ERPs and assessed schizotypy by the adjusted magical ideation scale (Eckblad and Chapman, 1983; Lustenberger et al., 2015). Thirty-five participants participated in this study. They were divided into two groups (low and high schizotypy) by a median split of the schizotypy scores. We found that the high-schizotypy group showed slowed mean reaction times in response to the face stimuli and the delay was correlated significantly with the schizotypy scores. In agreement with previous studies, we also found that the typical face-selective N170 component differed in amplitude between the face and non-face conditions and that the grand-averaged N170 component was significantly different over the occipito-temporal region. Between the two groups, importantly, we found reduced amplitude of the N170 components in the high-schizotypy group, which was significantly correlated with the schizotypy scores. Thus, our findings support the model that schizotypy is on a continuum with schizophrenia and that the neurophysiological substrate may represent where individuals are on this continuum.

The N170 has been considered a face-selective ERP component in healthy humans (Bentin et al., 1996; Rossion, 2014). However, one study claimed that this phenomenon reflects an artifact of uncontrolled inter-stimulus perceptual variance (ISPV) (Thierry et al., 2007). According to this study, a larger physical variance between stimuli caused an increased inter-trial jitter in the N170 for non-face stimuli thus the increased latency jitter reduced ERP amplitude. This study cast doubts on the well-established findings of face processing in human ERP studies. However, follow-up arguments claimed that there were methodological issues with the experimental images used in the study (Bentin et al., 2007; Rossion and Jacques, 2008). A further study also demonstrated that the N170 component is largely preserved after controlling for ISPV by presenting the same stimulus repeatedly (Ganis et al., 2012). In our study, we used a set of Mooney faces to minimize ISPV differences between the face and non-face conditions (Verhallen and Mollon, 2016) and investigated ISPV across the stimulus images in both face and non-face conditions. We computed progressive pixel-by-pixel averaging of the stimulus images in each condition (Figure 5A) and histogram of pixel-by-pixel correlations between the stimulus images for each condition to quantify ISPV (Figure 5B). We found no significant difference in mean correlations of the stimulus images between the face and non-face conditions (two-sample t-test, *p*>0.05). After controlling ISPV between the face and non-face conditions, we found larger amplitude of the N170 components over the occipito-temporal region for face stimuli compared to those for non-face stimuli (Figure 3). Thus, our findings of typical N170 components demonstrate that a face perception task using Mooney faces is an appropriate measure to investigate the face-selective N170 components.

**Figure 5.**
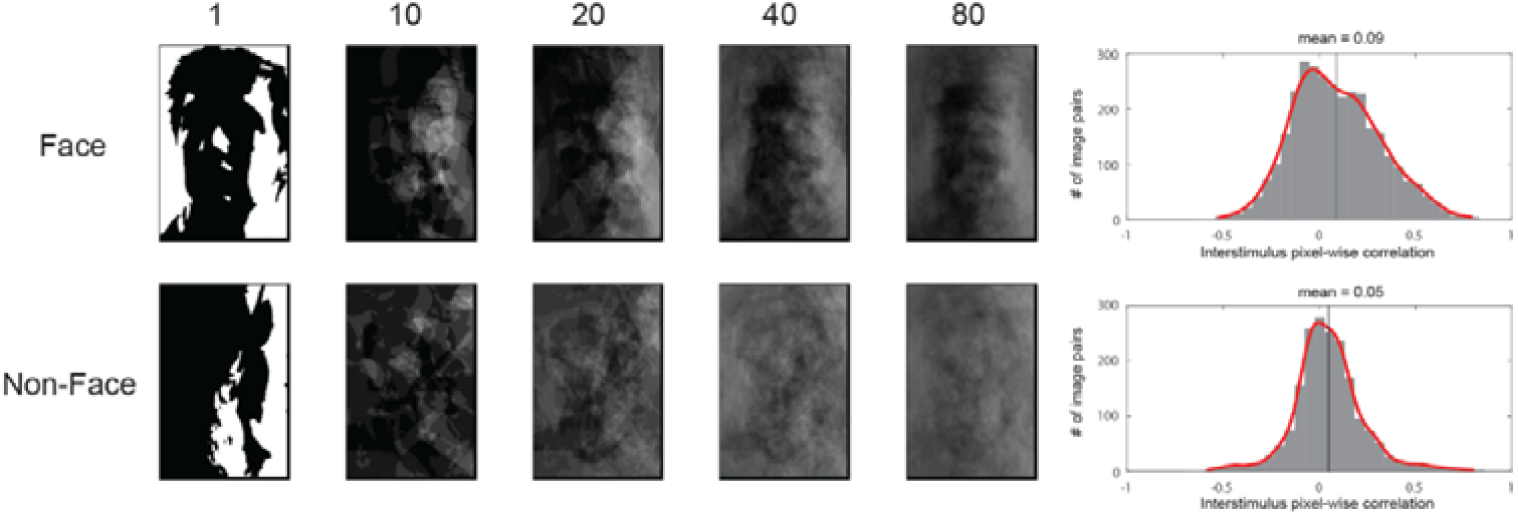
Inter-stimulus perceptual variance (ISPV) of stimulus images in face and non-face conditions. (A) Progressive pixel-by-pixel averaging of stimulus images in both conditions. The number of stimulus images used in each average is presented (B) Histogram of pixel-by-pixel correlations between stimulus images in each condition (Face: 0.09, Non-Face: 0.05). Red lines indicate fit lines to the histogram using non-parametric kernel-smoothing.

Impaired face perception has been reported in patients with schizophrenia (Herrmann et al., 2004; Kim et al., 2010; Shin et al., 2007; Soria Bauser et al., 2012). One study compared the N170 between patients with schizophrenia and healthy controls and found significantly lower differences in the N170 between face and building pictures in patients with schizophrenia (Herrmann et al., 2004). In addition to face perception, patients with schizophrenia responded more slowly and less accurately for body perception than healthy controls (Soria Bauser et al., 2012). Individuals at “ultra-high risk” for schizophrenia performed more poorly for face discrimination than healthy control (Kim et al., 2010). Importantly, we found slower reaction times 11 and reduced amplitudes of the N170 component in response to the face stimuli in the high-schizotypy group. Our findings support a theory of the psychosis continuum that similar symptoms that are observed in patients can be measured in non-clinical populations with higher incidence of schizophrenia-like experiences (van Os et al., 2009).

To date, few studies have investigated neurophysiological correlates in schizotypy (Aichert et al., 2012; Batty et al., 2014; Corlett and Fletcher, 2012). These investigations including our study are important to understand a continuum of psychosis disorders such as schizophrenia. One interesting finding that emerged was difference in reaction time but no difference in face discrimination accuracy (hit rate, Table. 2) between the low and high schizotypy groups. In contrast, previous studies reported that patients with schizophrenia exhibited lower face discrimination accuracy (Kim et al., 2010; Shin et al., 2007). These findings demonstrate that the processing of configural face perception in non-clinical populations who experience schizotypal traits may differ from impaired face recognition in patients with schizophrenia.

Our study has several limitations. First, we did not compare the N170 component with patients with schizophrenia. Although we found significant differences in behavioral and neurophysiological data between low and high schizotypy groups, age- and gender-controlled comparison is needed to investigate the proposed continuum of schizophrenia symptoms. A follow-up study should investigate both healthy controls and patients with schizophrenia for a direct comparison. Second, the schizotypy scores were not normally distributed (*p*<0.05, one-sample Kolmogorov-Smirnov test). However, the scores were well-distributed (i.e. little overlap across participants) thus it was assumed that the violation of normal distribution was not a critical factor for this study. Future studies should consider normality and administer additional measures for schizotypy such as the Schizotypal Personality Questionnaire (Raine, 1991).

In summary, we report the neurophysiological substrate of configural face perception in a non-clinical population who experience schizophrenia-like experiences such as magical ideation. We found significant differences in behavioral and neurophysiological data between the two groups and significant correlations with schizotypy scores. Our findings may support a theory of the psychosis continuum of schizophrenia and suggest that non-clinical populations may represent a useful target to investigate the neurobiological basis of schizophrenia not confounded by drug treatment.

## Acknowledgements

The authors thank Betsy Price, Jhana Parikh, and Jessica Page for helping data acquisition. The authors specially thank Roeland Verhallen for providing a set of Mooney face images. The authors gratefully acknowledge help and support from the Carolina Center for Neurostimulation.

## Contributions

C.L. and F.F. designed the study write the protocol. C.L. collected the data. S.A. analyzed the data. S.A. wrote the first draft of the manuscript. S.A., C.L., L.F.J., and F.F. edited the manuscript. All authors contributed to and have approved the final manuscript.

## Funding

This work was supported by the National Institute of Mental Health under Award Numbers R01MH111889 and R01MH101547.

## Conflict of Interest

S.A., C.L., L.F.J., and F.F. have no conflict of interest.

